# A molecular grammar for programmable multiphase protein–RNA vesicles

**DOI:** 10.64898/2026.03.04.709570

**Authors:** V. Ramachandran, D. A. Potoyan

## Abstract

Protein–RNA phase separation gives rise to biomolecular condensates with rich internal organization, yet the molecular rules that connect sequence-encoded interactions and composition to the emergent architecture of these condensates remain poorly defined. Here, using large-scale residue-level coarse-grained simulations, we identify a molecular grammar that governs the formation and stability of multiphase protein–RNA condensates. We show that asymmetries in protein–protein and protein–RNA interactions, together with protein stoichiometry, chain length, and condensate density, collectively determine whether condensates adopt homogeneous, layered, biphasic, or vesicle-like morphologies. Across a broad parameter space, these rules yield hollow multiphase vesicles with dense shells surrounding dilute interiors. Remarkably, vesicular condensates form spontaneously from well-mixed initial conditions, without requiring flux-driven oversaturation or extreme charge imbalance, distinguishing this mechanism from previously proposed routes to condensate hollowing. Our results establish minimal and general design principles for programming internal condensate architecture solely through sequence and composition, and provide a framework for engineering membrane-free vesicles and multilayered condensates with tunable permeability, encapsulation, and responsiveness.

## Introduction

Biomolecular condensates emerge from networks of weak, multivalent interactions that partition macromolecules into distinct mesoscale phases.^1–5^ By spatially and temporally organizing biochemical reactions, these condensates support a broad spectrum of cellular functions. Their assembly and dissolution are highly responsive to changes in physicochemical conditions, including temperature, pH, pressure, and acute environmental stress, enabling cells to sense perturbations, adapt internal organization, and maintain homeostasis.^6,7^ A notable example comes from yeast, where a starvation-induced drop in intracellular pH triggers the prion-like domain (PrD) of Sup35 to form nonfibrillar condensates.^8,9^

The internal organization of membraneless organelles is often far from uniform.^10,11^ Stress granules, for example, display a biphasic architecture consisting of a dense, solid-like core surrounded by a more dilute and dynamically exchanging shell.^12,13^ In contrast, *P* granules in *C. elegans* embryos exhibit a liquid core encased by a solid shell, arising from preferential localization of protein components to spatially distinct regions of the granule.^14^ When anchored to the nuclear pore, *C. elegans P* granules further assemble into a characteristic “tripartite sandwich” structure. ^15,16^ Such a multiphase organization and its ability to reorganize in response to cellular conditions underpin specialized functions and dynamic behaviors.

RNA–protein mixtures composed of low-complexity peptides and RNA have emerged as powerful *in vitro* model systems for probing the formation of multiphase condensates.^17–19^ In one example, RNA mixed with lysine and arginine forms droplets in which arginine preferentially associates with RNA to generate a dense core, while lysine is enriched at the periphery. Upon further addition of arginine, arginine displaces lysine from the interface, releasing lysine into the bulk.^20^ Similarly, in ternary mixtures containing a prion-like polypeptide (PLP), an arginine-rich polypeptide (RRP), and RNA, PLP and RNA compete for binding to RRP. This competition drives RNA-induced demixing, yielding coexisting phases comprising homotypic PLP-rich droplets and heterotypic RRP–RNA condensates. Depending on the RNA-to-RRP ratio and the relative strengths of underlying interactions, the resulting biphasic structures can adopt non-engulfing, partially engulfing, or fully engulfing morphologies.^18^ Together, these studies demonstrate that composition, preferential interactions, and stoichiometry jointly shape the architecture of multiphase condensates. Despite these advances, it remains unclear how sequence-encoded interactions and composition collectively select among distinct multiphase morphologies.

Although such behavior was historically associated with amphiphilic assemblies, recent studies have shown that RNA–protein mixtures can also form dynamic, membrane-free vesicles with well-defined hollow architectures under non-equilibrium conditions.^21,22^ However, a simple, general framework linking sequence-encoded interaction patterns and stoichiometry to the emergence of vesicular, layered, or biphasic condensates remains lacking. Here, we address this gap using a minimal coarse-grained model of protein–RNA condensates that captures essential sequence features while remaining computationally tractable. We design a low-complexity protein composed of two domains, H and L, with distinct propensities to interact with RNA and with each other (Fig. 1A), inspired by amphiphilic architectures that combine segments with contrasting affinities. By systematically tuning the relative strengths of Domain H–RNA, Domain L–RNA, and Domain L–L interactions, we define three classes of protein–RNA systems spanning regimes from weakly interacting auxiliary domains to strongly self-associating ones (Fig. 1B). Within each regime, we further vary the stoichiometric ratio of Domains H and L along the protein chain, while matching RNA length to the RNA-binding domain. Large-scale, residue-level coarse-grained simulations reveal a rich spectrum of condensate morphologies, including hollow multiphase vesicles, multilayered droplets with compositionally distinct shells, and biphasic condensates with dense cores (Fig. 1C). By analyzing domain-resolved radial distributions and effective interaction energies, we extract simple design rules that link molecular grammar, stoichiometry, density, and chain length to condensate architecture. Together, these results establish a mechanistic framework for understanding and engineering multiphasic protein–RNA condensates and generate concrete predictions for future reconstitution experiments.

**Figure 1:**
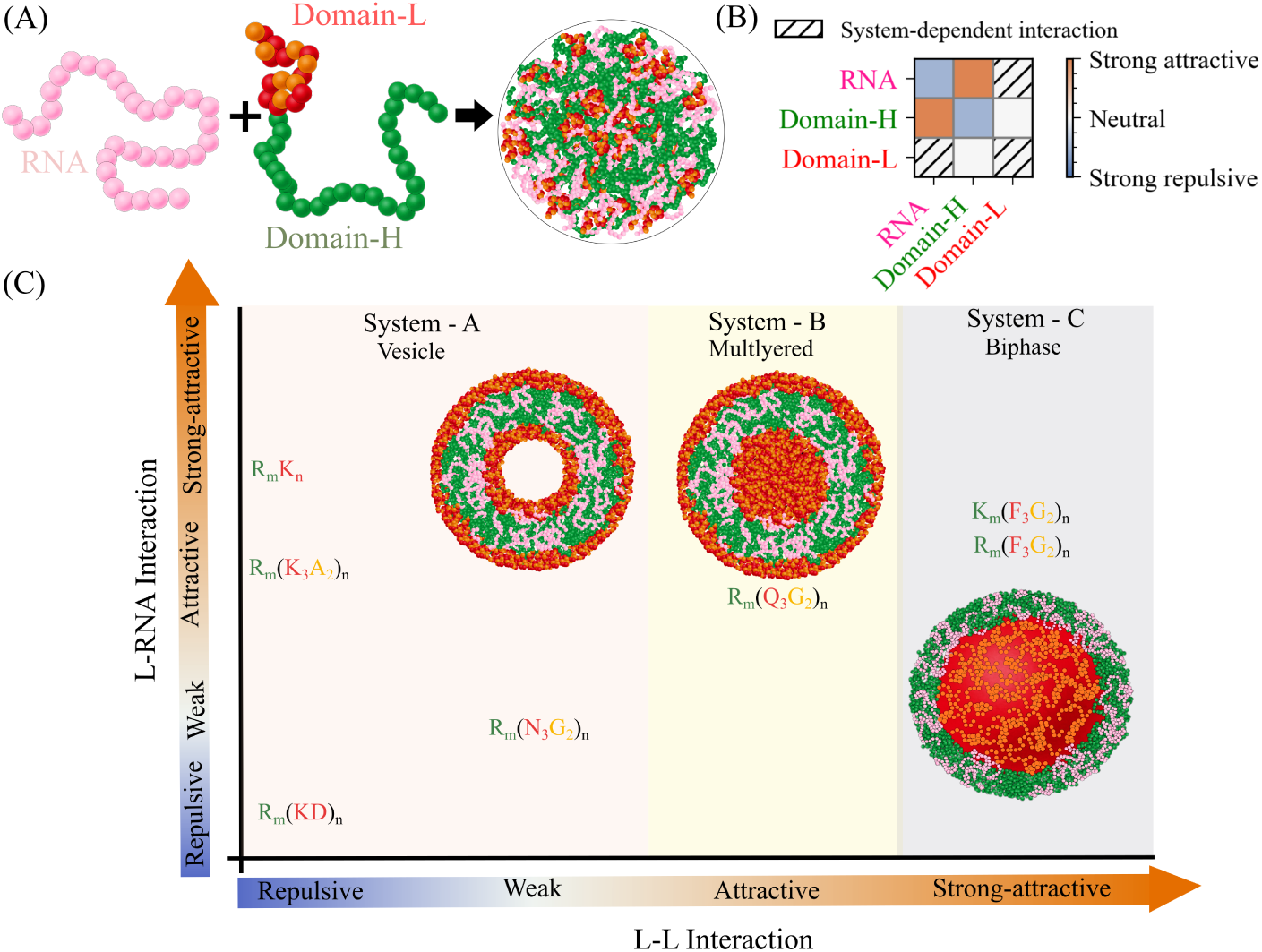
(A) Schematic representation of the simulation setup. (B) Interaction matrix of domains (C) Sequence space of the condensates simulated as a function of Domain L–L and L–RNA interactions.

## Methods

### Residue-Resolution Coarse-Grained Model

We employ a residue-level coarse-grained (CG) model to study protein–RNA condensates.^23,24^ In this model, a harmonic potential is used for bonded interactions between adjacent beads,

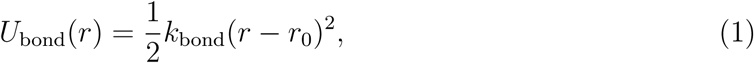

with a bond force constant *k*_bond_ = 1000 kJ mol*^−^*^1^ nm*^−^*^2^. The equilibrium bond length is set to *r*_0_ = 0.5 nm for RNA and *r*_0_ = 0.38 nm for peptides.^25^

Non-bonded interactions are modeled using the Ashbaugh–Hatch (AH) potential for short-range interactions, combined with a Debye–Hückel (DH) potential to account for screened electrostatics. The AH potential is given by

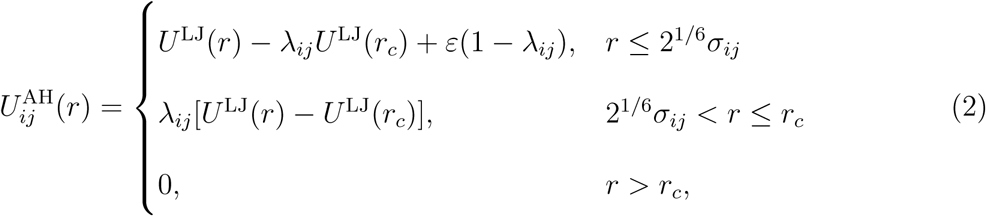

where *U* ^LJ^(*r*) is the Lennard–Jones potential, *σ_ij_* = (*σ_i_*+*σ_j_*)*/*2 is the arithmetic mean of bead diameters, and *λ_ij_* = (*λ_i_* + *λ_j_*)*/*2 is the mean hydrophobicity parameter. When *λ* = 0, the AH potential reduces to a purely repulsive Weeks–Chandler–Andersen (WCA) interaction.^26^ Amino acid bead diameters *σ_i_* are assigned based on van der Waals radii.^27^ At the same time, all nucleotides are represented by a single bead type with fixed diameter *σ* = 0.817 nm and hydrophobicity *λ* = −0.027. Protein hydrophobicity parameters are taken from the CALVADOS 2 force field.^24^

Electrostatic interactions are modeled using the Debye–Hückel potential,

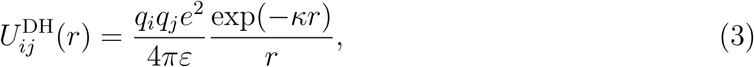

where *q_i_* and *q_j_* are the bead charges. The inverse Debye screening length *κ* is determined from the ionic strength *I* as

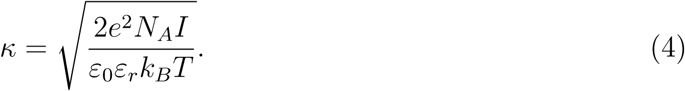

All simulations were performed at ionic strength *I* = 0.1 nm*^−^*^3^, corresponding to a screening length *κ^−^*^1^ = 0.96 nm at *T* = 300 K. The temperature dependence of the dielectric constant was incorporated using the empirical relation

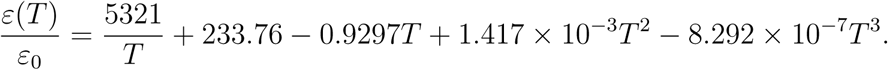

Simulations were carried out using a time step of 10 fs and continued until stable condensates formed. Systems were simulated in cubic boxes with linear dimensions between 120 and 160 nm, containing 900–1600 chains per component. A complete summary of simulation parameters is provided in Table S2.

### Analysis

To characterize the spatial organization of proteins and RNA within condensates, we computed radial distribution functions (RDFs) relative to the condensate center of mass. The potential of mean force (PMF) was obtained from the RDF *g*(*r*) via

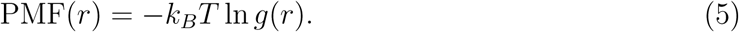

To obtain a single representative measure of effective interaction strength, the PMF was averaged over distance using a probability-weighted mean, consistent with recent sequencebased frameworks that leverage coarse-grained energy functions to define effective interaction energies between disordered regions.^28^

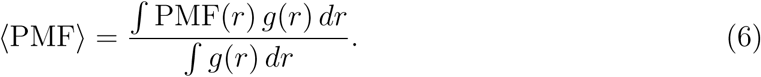

This quantity captures the dominant contribution to pairwise interactions within the condensate. Local interaction patterns were further quantified by computing the average number of RNA or protein neighbors within a cutoff distance of 11 Å around each protein residue. These contact numbers were averaged over all residues and chains, providing a direct measure of local protein–RNA and protein–protein interaction strengths across different systems. All other structural analyses have been carried out using the MDAnalysis package.^29^

## Results

### Protein Sequence Grammar Dictates Multiphase Condensate Architecture

Protein composition plays a central role in determining the internal organization of multiphase condensates.^10,20,30–33^ For instance, mixtures of two distinct proteins and RNA can give rise to rich multiphase architectures as a function of protein composition.^18,34,35^ To dissect how specific protein–RNA and protein–protein interactions encode the emergent multi-phase architectures of condensates, we engineer a low-complexity protein composed of two distinct regions, domains H (high RNA affinity) and L (low RNA affinity). While both domain composition and stoichiometry influence multiphasic morphology, here we focus exclusively on the role of protein grammar, defined by the amino acid composition of Domains H and L.

By design, Domain H-L interactions are kept comparatively weak, with a strong attractive interaction of Domain H-RNA; a detailed interaction matrix of the domain is shown in Fig. 1B. Within this framework, we then systematically tune Domain L-RNA and L-L interaction strengths by altering the amino acid composition of Domain L (Fig. 1C). This strategy allows us to isolate how changes in Domain L chemistry control condensate morphology and internal organization.

Based on these interaction patterns, we define three classes of protein–RNA systems. System A is characterized by weak Domain L-L interactions, with Domain L–RNA interactions ranging from repulsive to strongly attractive (Fig. 1C). System B is characterized by attractive Domain L-L interactions combined with weak Domain L–RNA affinity (Fig. 1C). System C is characterized by strong L-L interactions with Domain L–RNA interactions that are weaker than those of Domain H–RNA (Fig. 1C). For each system, we select amino acid compositions for Domain H and L that realize the targeted interaction profiles (Table S1). We then perform long-timescale molecular dynamics simulations using the CALVADOS-2 single-bead coarse-grained model.^24^ Simulation details, including chain numbers and ionic conditions, are summarized in Table S2.

In our notation, protein sequences are written as *A_m_B_n_*, where the first term *A_m_* denotes Domain H composed of *m* residues of amino acid A, and the second term *B_n_* denotes Domain L composed of *n* residues of amino acid B. In System A, Domain L-L interactions are designed to be repulsive or weak, while Domain L–RNA interactions span a range from repulsive to weak to attractive (Fig. 1B). To realize this interaction landscape, we construct four protein compositions: (i) *R*_150_(*KD*)_25_, in which both Domain L-L and Domain L–RNA interactions are repulsive due to the presence of aspartic acid (D); (ii) *R*_150_(*K*_3_*A*_2_)_10_, where Domain L-L interactions remain electrostatically repulsive because of lysine (K), while Domain L–RNA interactions become attractive due to alanine (A); (iii) *R*_150_*K*_50_, which exhibits repulsive Domain L-L interactions and attractive Domain L–RNA interactions, albeit weaker than those of Domain H–RNA, such that RNA preferentially associates with Domain H; and (iv) *R*_150_(*N*_3_*G*_2_)_10_, in which both Domain L-L and Domain L–RNA interactions are uniformly weak.

Across all four compositions in System A, we observe the robust formation of multiphasic vesicles. These vesicles exhibit a characteristic architecture consisting of an empty core, an inner shell enriched in Domain L, a middle shell composed of Domain H–RNA complexes, and an outer Domain L–rich shell (Fig. 2B–F). This layered organization is directly reflected in the radial distribution functions (RDFs) of the domains relative to the condensate center of mass (Fig. 2B–E). To our knowledge, this specific multiphase vesicle architecture has not been reported previously. In this configuration, Domain L segments, together with Domain H–RNA complexes, effectively behave as composite monomers analogous to amphiphilic molecules, whose segregated chemical functionalities drive liposome formation^36^ (Fig. 2A).

**Figure 2:**
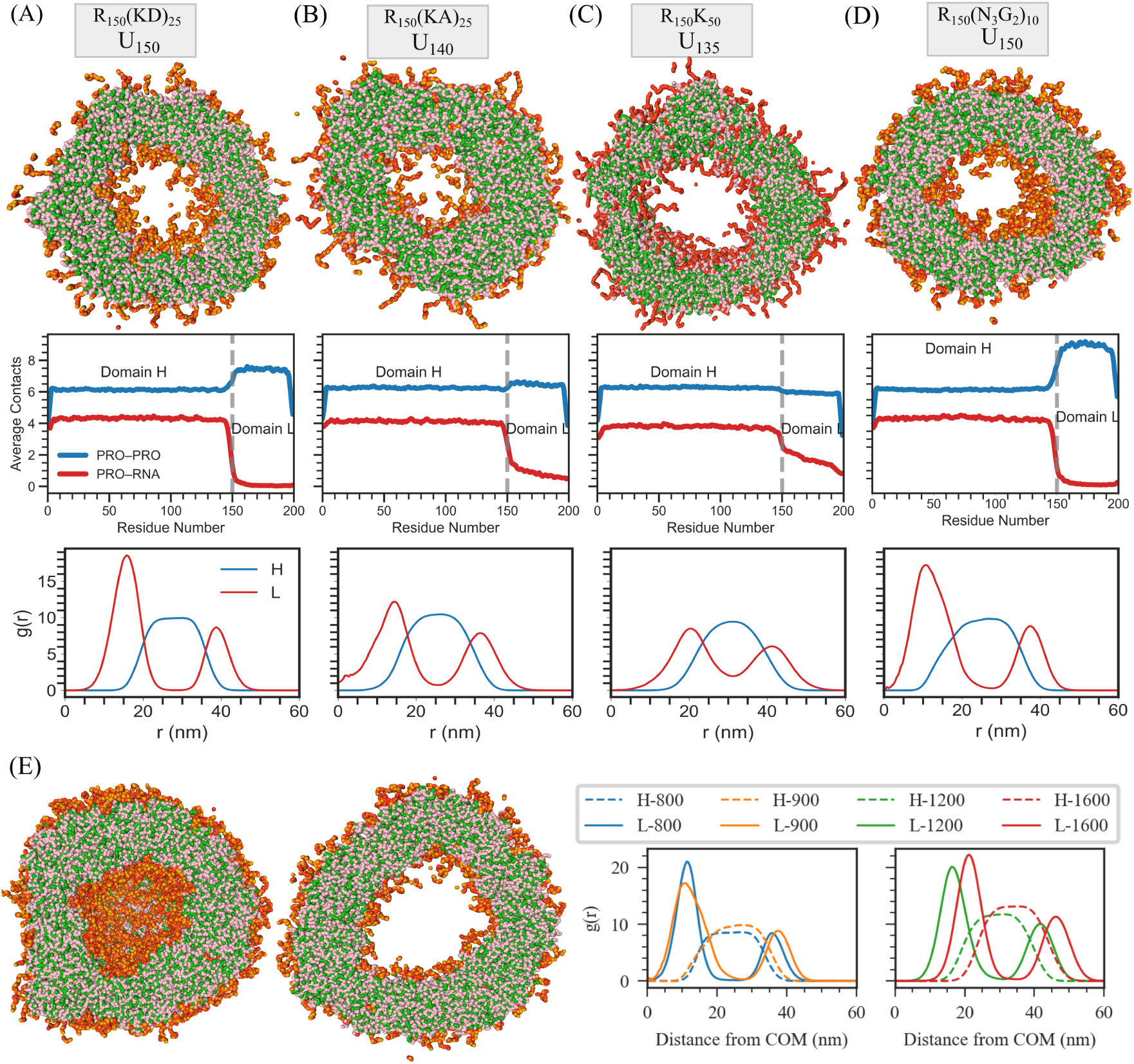
Structural characterization of multiphase vesicles in System A. (A–D) Top: Vesicle slice snapshots; middle: Corresponding average number of contacts surrounding the protein, including protein–protein (denoted as PRO) and protein–RNA contacts; bottom: Corresponding radial distribution functions (RDFs) of Domain L and Domain H–RNA complexes around the condensate center of mass for four sequences: (A) R_150_(KD)_25_ with U_150_, (B) R_150_(K_3_A_2_)_10_ with U_140_, (C) R_150_K_50_ with U_135_, and (D) R_150_(N_3_G_2_)_10_ with U_150_. (E) Size dependence of vesicle formation for the R_150_(N_3_G_2_)_10_ with U_150_ system. Left: snapshot of a vesicle formed by 1600 chains of each component, showing a pronounced hollow core. Right: RDF around the center of mass (COM) of the condensate as a function of the number of chains, demonstrating that the core size increases with system size. Color scheme: Green denotes arginine (R) of Domain H, pink denotes uracil (U), and red represents lysine (K) or asparagine (N), and orange represents glycine (G) or alanine (A), collectively forming Domain L.

Vesicle formation is primarily driven by the preferential affinity of RNA for Domain H (arginine-rich) over Domain L, combined with weak and nonspecific interactions within Domain L. This interaction hierarchy is quantified by computing the average number of neighboring RNA or protein residues within 11 Å of each protein residue, which clearly differentiates domain-specific protein–protein and protein–RNA interaction patterns across compositions spanning repulsive to attractive regimes (Fig. 2B–E).

To assess the robustness of vesicle formation, we further simulate the *R*_150_(*N*_3_*G*_2_)_10_ system across a range of system sizes by varying the number of chains from 800 to 1600 (corresponding to 281,600–563,200 particles) and adjusting the simulation box dimensions accordingly (Fig. 2F; Fig. S1; Table S2). In all cases, stable vesicles form, with the size of the empty core increasing systematically with system size (Fig. 2F). Due to the substantial computational cost, system-size scaling was examined only for this representative composition.

RNA concentration plays a decisive role in shaping condensate morphology, particularly when Domain L interacts attractively with RNA. At *U*_150_, the *R*_150_(*KD*)_25_ and *R*_150_(*N*_3_*G*_2_)_10_ systems readily form multiphasic vesicles, whereas the *R*_150_(*K*_3_*A*_2_)_10_ and *R*_150_*K*_50_ systems produce non-vesicular condensates under identical conditions (Fig. 3; Fig. S2). Strikingly, reducing the RNA content to *U*_135_ for *R*_150_(*K*_3_*A*_2_)_10_ and to *U*_140_ for *R*_150_*K*_50_ induces vesicle formation in both cases (Fig. 3; Fig. S2). These results indicate that when Domain L–RNA interactions are attractive, limiting RNA availability enhances competition between protein–protein and protein–RNA interactions, thereby promoting vesicular architectures.

**Figure 3:**
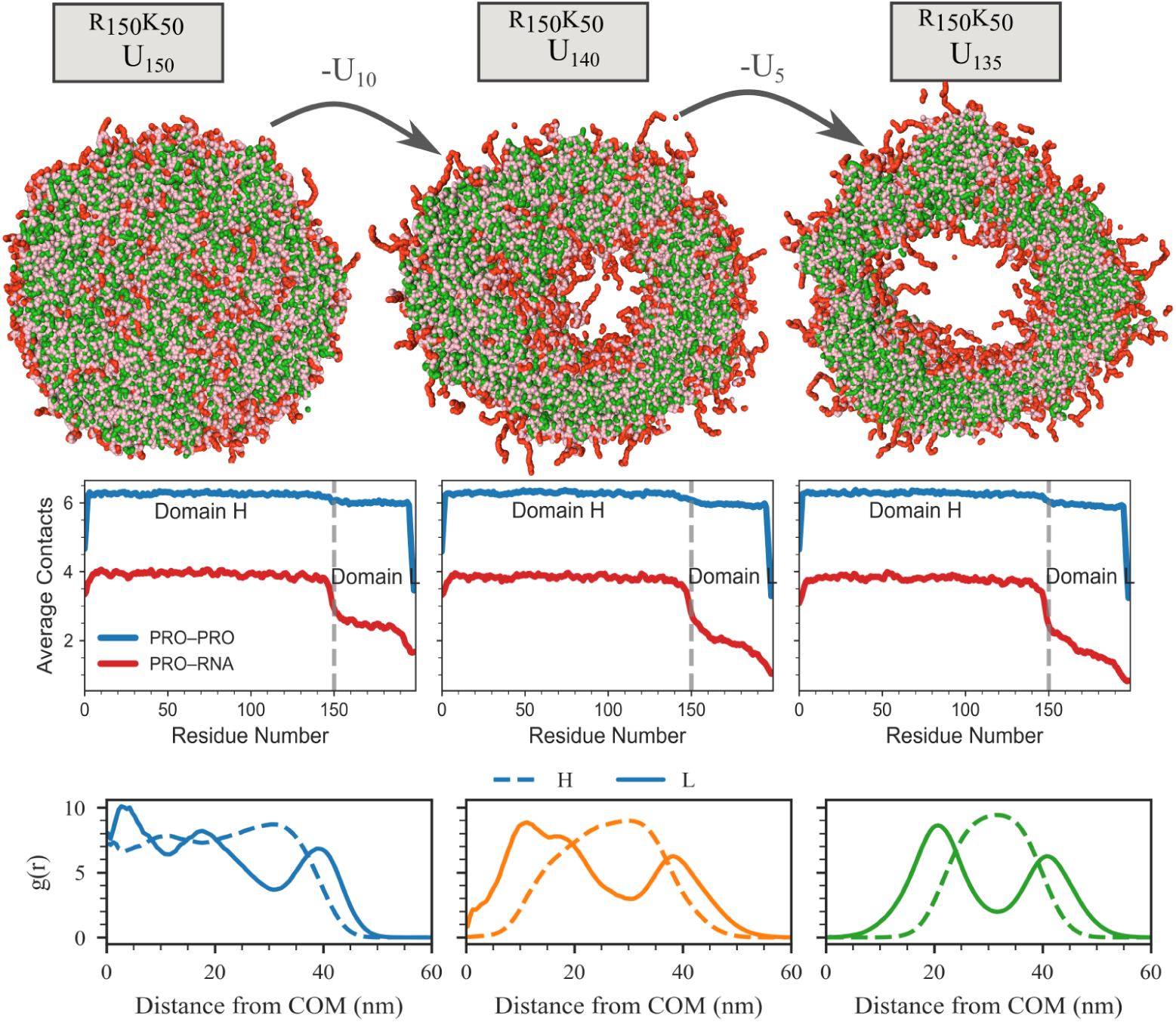
Influence of RNA concentration on vesicle formation in the R_150_K_50_ sequence. Top: Condensate snapshots of R_150_K_50_ with U_150_, U_140_, and U_135_. Middle: Corresponding average number of contacts surrounding the protein components. Bottom: Radial distribution functions (RDFs) around the center of mass (COM) for Domain L and Domain H–RNA complexes.

Consistent with this interpretation, contact analysis at *U*_150_ reveals diminished preferential binding of Domain H to RNA relative to Domain L in the *R*_150_(*K*_3_*A*_2_)_10_ and *R*_150_*K*_50_ systems. Upon RNA reduction, strong Domain H–RNA preference is restored (Fig. 3). Together, these trends show that within System A, vesicle formation is driven either by strong repulsion in both Domain L-L and Domain L–RNA interactions or by attractive Domain L–RNA interactions under conditions of limited RNA availability.

To transition from System A to System B, we introduce attractive Domain L-L interactions while maintaining weak Domain L–RNA affinity. Specifically, we replace asparagine (N) with glutamine (Q) in Domain L, yielding the composition *R*_60_(*Q*_3_*G*_2_)_28_, which promotes favorable Domain L-L contacts. Contact analysis confirms the emergence of attractive Domain L-L interactions, as reflected by an increased number of neighboring Domain L residues within 11 Å of each protein (Fig. 3).

In System B, we observe a distinct, layered, multiphase architecture that differs qualitatively from the hollow vesicles formed in System A. The condensate consists of a Domain L–rich core, surrounded by a shell of Domain H–RNA complexes, and further enveloped by an outer Domain L–rich layer, as revealed by the radial distribution functions (RDFs) relative to the condensate center of mass (Fig. 4A). Notably, the central core exhibits a lower density than the surrounding layers, consistent with a filled, loosely packed interior rather than a hollow lumen (Fig. 4A). These observations indicate that attractive Domain L-L interactions drive core formation, fundamentally altering the condensate architecture relative to System A.

**Figure 4:**
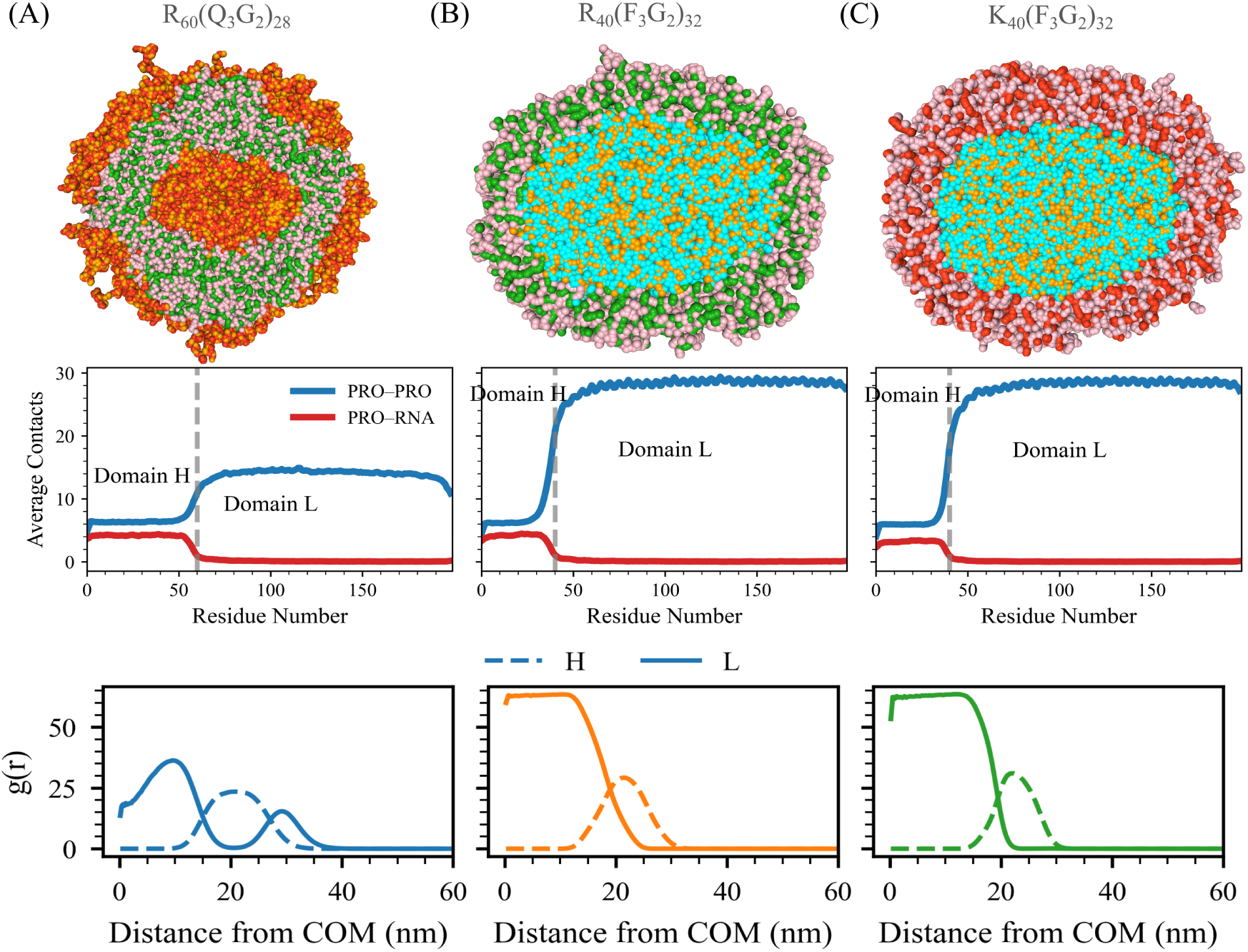
Structural characterization of multiphase condensates in Systems B and C. (A–C) Top: Cross-sections of the condensates; middle: corresponding average number of contacts surrounding the protein components; bottom: corresponding radial distribution functions (RDFs) of Domain L and Domain H–RNA complexes around the condensate center of mass for System B and System C: (A) System B; R_60_(Q_3_G_2_)_28_ with U_60_, (B) System C; R_40_(F_3_G_2_)_32_ with U_40_, and (C) System C; R_40_(F_3_G_2_)_32_ with U_40_. Color scheme: green denotes arginine (R), pink denotes uracil (U), red represents glutamine (Q) or lysine (K), cyan represents phenylalanine (F), and orange represents glycine (G).

In System C, where Domain L-L interactions are strongly attractive and Domain L–RNA interactions are also attractive but weaker than Domain H–RNA affinity, we examine two compositions: *K*_40_(*F*_3_*G*_2_)_32_ and *R*_40_(*F*_3_*G*_2_)_32_. Within the CALVADOS-2 model, phenylalanine–phenylalanine interactions are strongly attractive, as reflected by elevated numbers of neighboring Domain L residues within 11 Å of each protein (Fig. 4B–C). In both systems, this interaction hierarchy produces biphasic condensates composed of a dense Domain L–rich core surrounded by an outer shell of Domain H (lysine- or arginine-rich)–RNA complexes, as evidenced by their RDF profiles (Fig. 4B–C).

Taken together, systematic tuning of competitive protein–protein and protein–RNA interactions generates a diverse spectrum of multiphase condensate architectures, spanning hollow vesicles, multilayered droplets, and biphasic condensates.

### Stoichiometry of Protein Governs Multiphase Condensate Formation

Nature of the condensates from uniform droplets to multiphase condensates and hollow vesicles is driven not only by the “protein grammar of domains” that encodes interaction motifs, but also by the stoichiometric ratio of Domains H and L in a protein. To examine this, we systematically vary the relative ratio of Domains H and L in 200-residue-long proteins, while keeping the RNA length precisely matched to the length of Domain H, across all three model systems (Systems A, B, and C).

Starting with System A, we use the baseline composition R*_m_*(N_3_G_2_)*_n_*with U*_m_*, in which both Domain L–L and Domain L–RNA interactions are weak. We systematically vary the parameters *m* and *n*, thereby generating nine distinct protein sequences (Fig. 5). In all cases, the RNA length is chosen to match the number of arginine residues in Domain H.

**Figure 5:**
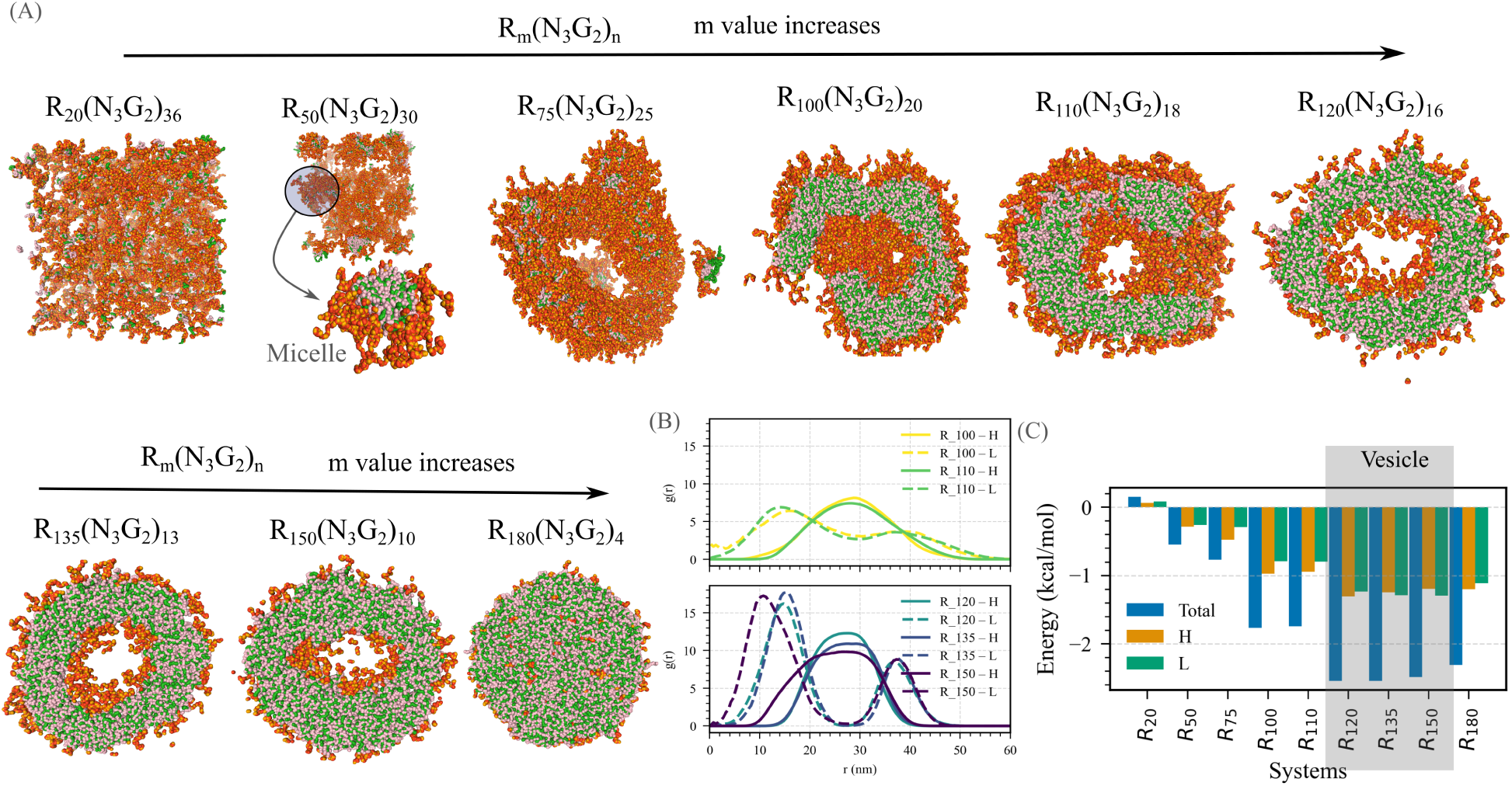
Stoichiometric evolution of condensate morphology in System A. (A) Representative simulation snapshots illustrating how condensate structure changes with varying ratios of Domain H (R*_m_*) to Domain L ((N_3_G_2_)*_n_*). The sequences (from left to right) correspond to R*_m_*(N_3_G_2_)*_n_* with (*m, n*) values of (20, 36), (50, 50), (75, 25), (100, 20), (110, 18), (120, 16), (135, 13), (150, 10), and (180, 4). (B) Radial distribution functions (RDFs) of Domain L and Domain H–RNA around the condensate center of mass, comparing partially formed vesicle sequences (top) and fully developed vesicle sequences (bottom). (C) Potential of mean force (PMF) energy weighted by the RDF as a function of the Domain H–to–Domain L ratio, highlighting the energetic trends underlying vesicle formation. Color scheme: green denotes arginine (R) (Domain H), pink denotes uracil (U), red represents asparagine (N), and orange represents glycine (G), collectively forming Domain L.

Across this sequence space, we observe a clear, stoichiometry-driven evolution of condensate morphology. The internal organization of condensates changes systematically with *m* (or with the relative ratio of Domain H to Domain L in a sequence). Sequences with the shortest Domain H (or lowest *m*, corresponding to the longest relative Domain L segment) fail to undergo condensation, consistent with the lack of sufficient favorable interactions needed for assembly (Fig. 5A).

As the ratio of Domain H (R*_m_*) in a sequence increases, it begins to form multiple micelle-like droplets, characterized by a Domain H–RNA-rich core surrounded by a Domain L ((N_3_G_2_)*_n_*) shell (Fig. 5A). Further increasing the relative ratio of Domain H in a sequence produces doughnut-shaped condensates, which preserve the same core–shell organization. Pushing Domain H ratio in a sequence even further results in the emergence of partially formed vesicles containing multiple openings, a morphology that, to our knowledge, has not previously been reported (Fig. 5A). Radial distribution functions of individual Domains around the center of mass of condensate confirm this progression, revealing incomplete, multiphase vesicle-like structures in the Domain H-RNA regime (Fig. 5B).

With a further increase of Domain H ratio in a sequence, it transitions into fully developed multiphasic vesicles. These vesicles feature a hollow core, an inner shell enriched in Domain L, a middle layer composed of Domain H–RNA complexes, and an outer Domain L shell. This multilayered architecture is clearly reflected in the radial distribution functions, which show well-resolved, concentric domain-specific layers surrounding an empty core (Fig. 5B).

At the highest Domain H ratios in a sequence, however, the vesicular structure is lost. Instead, it forms a multiphase condensate with a Domain H–RNA–rich interior surrounded by the Domain L-domain ((N_3_G_2_)*_n_*) layer at the periphery (Fig. 5A).

We compute the potential of mean force (PMF) energy weighted by the RDF, to obtain a single scalar measure that reflects the overall effective interaction between the species and thereby determines structural stability. A more negative value of this energy indicates stronger attractive interactions and a greater tendency to form. As shown in Fig. 5C, PMF energy varies systematically with the Domain H–to–Domain L ratio. Notably, the energy becomes highly negative within the vesicle-forming stoichiometric window, consistent with strong inter-Domain H attraction driving vesicle assembly. In contrast, the energy remains positive for sequences with the highest Domain-L ratio in the sequence, where no clustering or vesicle formation is observed.

Overall, altering the stoichiometry of Domains H and L of System A drives a remarkable range of architectures, from non-condensed states to micelle-like droplets, partially closed vesicles, fully multiphase vesicles, and ultimately to multiphase condensates.

Next, in System B, where Domain L–L, Domain L–RNA, and Domain H–RNA interactions are all attractive, the baseline composition is R*_m_*(Q_3_G_2_)*_n_*with U*_m_*. We generate six protein sequences by systematically varying *m* and *n* (Fig. 6A). In contrast to System A, stable condensates form across all stoichiometries due to the presence of multiple favorable interactions.

**Figure 6:**
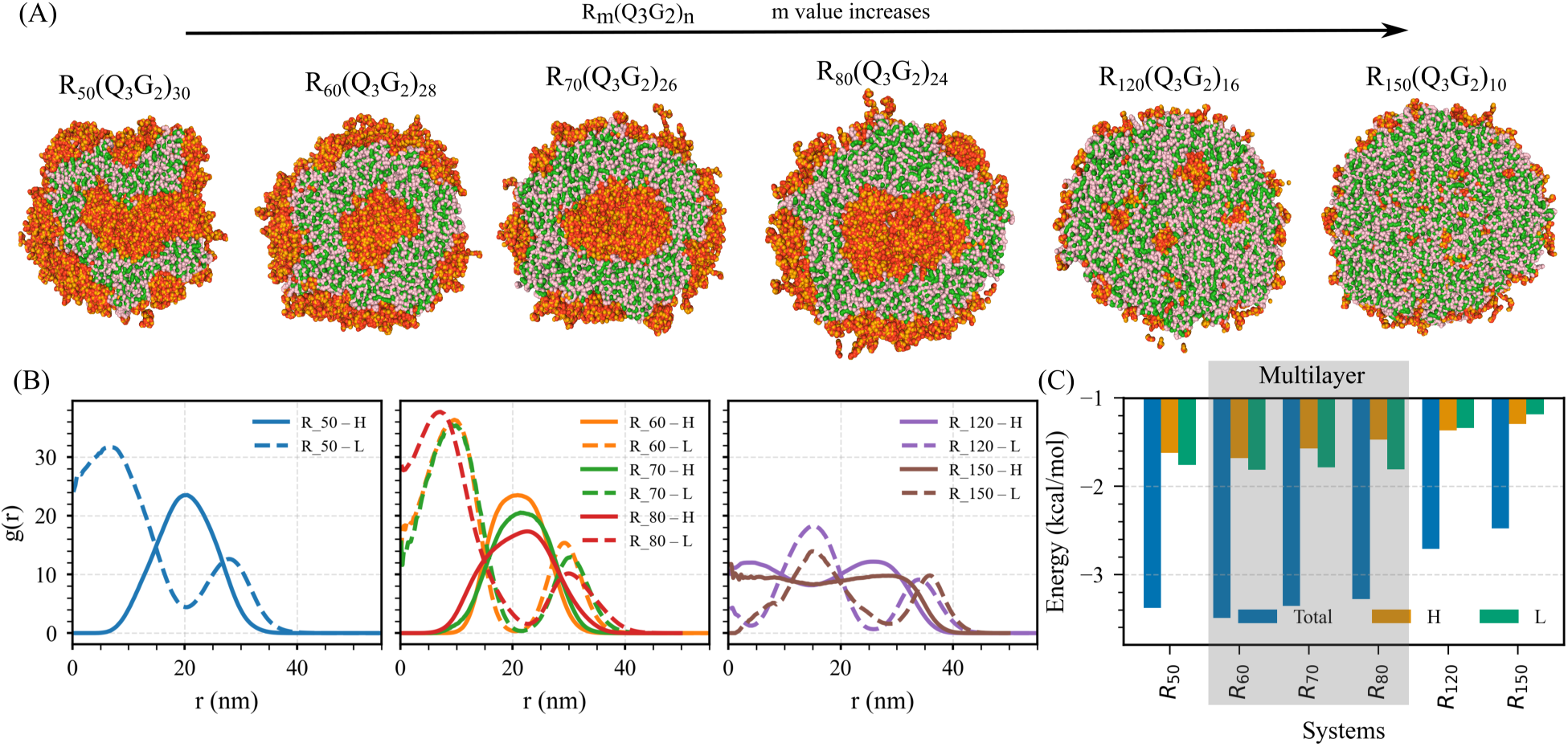
Stoichiometric evolution of condensate morphology in System B. (A) Representative simulation snapshots showing condensate evolution with the ratio of Domain H (R*_m_*) to Domain L ((Q_3_G_2_)*_n_*) in System B. The sequences shown (from left to right) correspond to R*_m_*(Q_3_G_2_)*_n_* with (*m, n*) values of (50, 30), (60, 28), (120, 16), (135, 13), and (150, 10). (B) Radial distribution functions (RDFs) of Domain L and Domain H–RNA complexes around the condensate center of mass. (C) Potential of mean force (PMF) energy weighted by the RDF as a function of the Domain H–to–Domain L ratio, highlighting the energetic trends underlying layered structure formation. Color scheme: green denotes arginine (R) of Domain H, pink denotes uracil (U), red represents asparagine (N), and orange represents glycine (G), collectively forming Domain L.

The internal organization of these condensates evolves systematically with increasing *m* (or with the relative ratio of Domain H to Domain L in a sequence). At a low Domain H ratio in a sequence, the condensates adopt a multilayered architecture consisting of a Domain L-rich core, a Domain H–RNA-enriched intermediate layer, and an outer shell enriched with Domain L ((Q_3_G_2_)*_n_*) (Fig. 6A). Importantly, the intermediate Domain H–RNA layer does not fully encapsulate the Domain L–rich core, as reflected in the radial distribution functions (Fig. 6A,B).

As the Domain H (R*_m_*) ratio increases in a sequence, the intermediate Domain H–RNA layer becomes thicker, more continuous, and more compositionally enriched, ultimately forming a fully developed multilayered structure in which the Domain H–RNA layer entirely covers the Domain L-rich interior (Fig. 6A,B). With further increases of the Domain H ratio in a sequence, the Domain L-rich regions begin to fragment into smaller, irregular clusters embedded within a continuous Domain H–RNA matrix. At the same time, a thin Domain L-rich rim persists at the periphery of the condensate (Fig. 6A).

In the highest Domain H ratio in sequences we have simulated, Domain L-rich were greatly diminished, appearing only as small puncta dispersed throughout the Domain H– RNA network, with the outer surface retaining only a faint enrichment of Domain L (Fig. 6A,B).

Similar to System A, the PMF energy weighted by RDF was computed as a function of the Domain H–to–Domain L ratio. Unlike System A, the PMF energy remains negative across all compositions, indicating overall attractive interactions and stable condensate formation. Notably, the multilayered vesicle-forming compositions exhibit the most negative PMF energy, reflecting highly favorable interaction patterns within multilayered structures (Fig. 6C).

Overall, increasing the Domain H ratio in sequences drives the condensates from poorly defined multilayer structures to well-defined multilayers, scattered Domain L-rich domains in the Domain H–RNA complex, and finally to a biphasic structure.

Finally, in System C of a baseline composition of R*_m_*(F_3_G_2_)*_n_*with U*_m_*, we have constructed five protein sequences by systematically varying *m* and *n* (Fig. 7A). As in System B, Domain L–L, Domain L–RNA, and Domain H–RNA interactions are attractive, enabling condensate formation across all stoichiometries.

**Figure 7:**
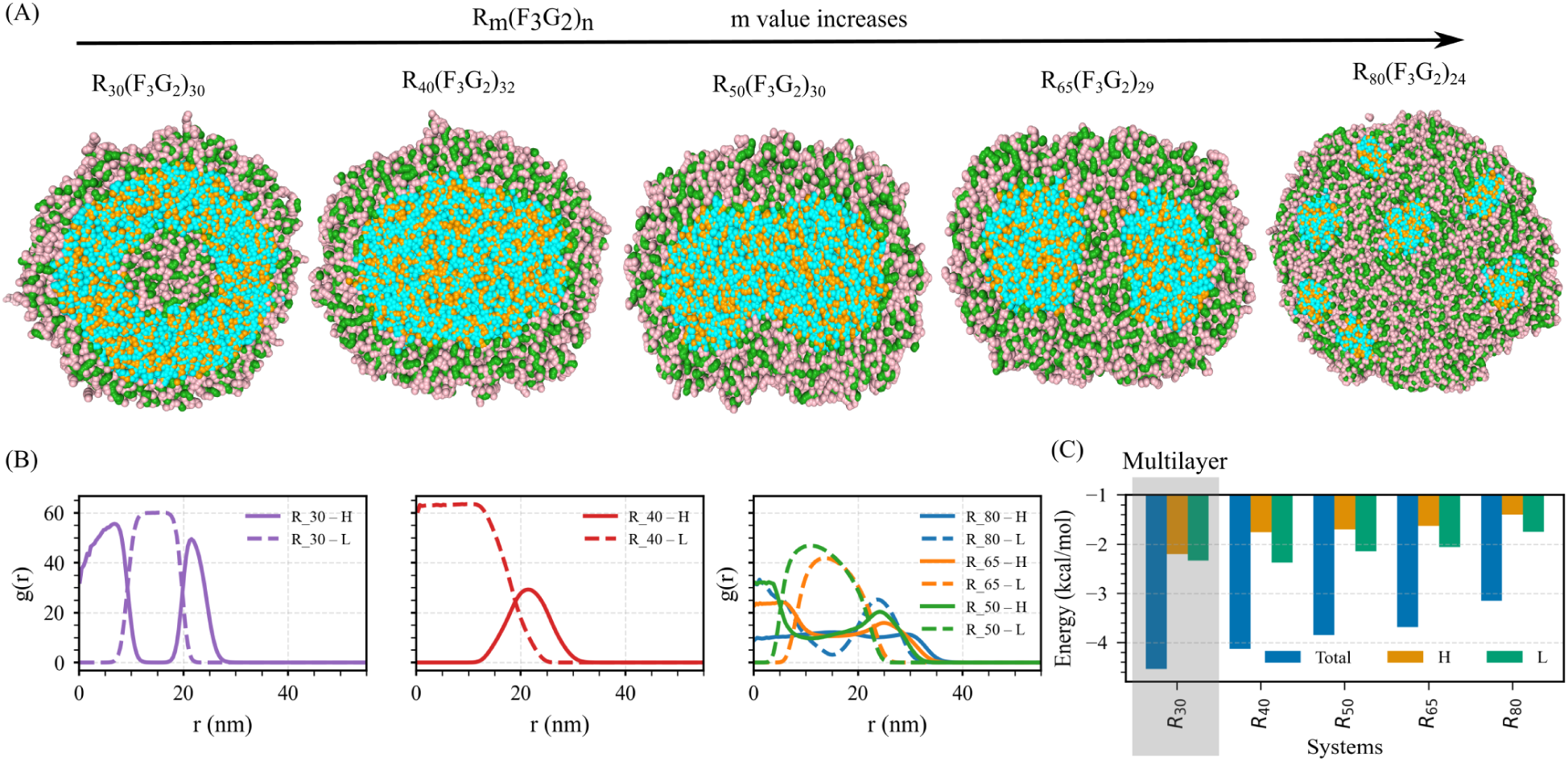
Stoichiometric evolution of condensate morphology in System C. (A) Representative simulation snapshots illustrating condensate morphology at different stoichiometries. The sequences shown (from left to right) correspond to R*_m_*(F_3_G_2_)*_n_* with (*m, n*) values of (30, 30), (40, 32), (50, 30), (65, 29), and (80, 24). (B) Radial distribution functions (RDFs) of Domain L and Domain H–RNA around the condensate center of mass, demonstrating progressive changes in internal structure with increasing Domain H content. (C) Mean potential of mean force (PMF) energy weighted by the RDF as a function of the Domain H– to–Domain L ratio, highlighting the energetic trends underlying layered structure formation. Color scheme: green denotes arginine (R), pink denotes uracil (U), red represents glutamine (Q) or lysine (K), cyan represents phenylalanine (F), and orange represents glycine (G), collectively forming Domain L.

At the lowest Domain-H ratio of a sequence, the condensates adopt a multilayered architecture consisting of a Domain H–RNA-rich core, surrounded by a Domain L intermediate layer, and capped by an outer Domain H–RNA-rich shell (Fig. 7A and 7B). This arrangement resembles the multilayer organization observed in System B, but with the layering order reversed.

As the ratio of Domain H in a sequence increases, it transitions to a biphasic condensate characterized by a Domain L-rich core enveloped by a continuous Domain H–RNA shell, a structure clearly reflected in the radial distribution functions (Fig. 7A,B). Further enrichment of Domain H leads to the penetration of Domain H–RNA complexes into the central Domain L-rich region, producing a partially mixed core (Fig. 7A,B).

At the highest Domain H ratio in a sequence examined, the condensates become predominantly Domain H–RNA rich, with only small, isolated Domain L-rich clusters dispersed within the continuous Domain H–RNA matrix (Fig. 7A). This progression illustrates how increasing the abundance of Domain H gradually transforms the condensate from a Domain L-centered, layered architecture into one dominated by Domain H–RNA interactions, with Doain L-rich components reduced to scattered inclusions.

Similar to Systems A and B, the PMF energy weighted by RDF was computed as a function of the Domain H–to–Domain L ratio. Similar to System B, the PMF energy remains negative across all compositions, indicating overall attractive interactions and condensate stability. Notably, the multilayered vesicle-forming compositions exhibit the most negative mean PMF energy, reflecting highly favorable interaction patterns within multilayered structures (Fig. 7C).

Overall, both protein grammar and Domain H:L stoichiometry act in concert to govern condensate morphology, enabling the fine-tuning of multiphase architectures.

### Influence of density and chain length on multiphase vesicle formation

The density and chain length of biomolecules play pivotal roles in governing both the kinetics and thermodynamics of liquid–liquid phase separation (LLPS).^37–42^ Usually, LLPS is only favorable once the biomolecular chain length and concentration exceed critical thresholds.^43^ In simulations, the density of the system is chosen pragmatically: below the typical liquid phase density (∼500 mg/mL) but well above the solution phase density used in experiments (∼1 mg/mL). Higher density ensures rapid droplet formation (*<* 500 ns) via diffusion and binding mechanisms, thereby facilitating the sampling of different droplet compositions within reasonable computational time.

To systematically evaluate the influence of biomolecular density on the kinetics and thermodynamics of RNA–protein LLPS, and additionally to test the robustness of our findings, we have performed simulations at multiple densities using a small model system R_135_(N_3_G_2_)_13_ with U_135_ consisting of 800 chains each, for a total of 1600 chains. They form a vesicle-like condensate. We employ a uniform simulation protocol adapted from methodologies widely used in lipid vesicle molecular dynamics simulations.^44,45,45,46^ First, all biomolecules are uniformly distributed in the simulation box using Packmol.^47^ To further homogenize the initial configuration, we perform a brief simulation in which repulsive forces are applied to prevent premature clustering and ensure a well-mixed starting state. Following this, the simulation box is expanded by 25 nm in all spatial dimensions. This enlargement reduces the likelihood that early aggregates interact with their periodic images, a key precaution to avoid unphysical fusion events that could otherwise promote the formation of lamellar phases rather than closed vesicle structures.

We have constructed nine simulation systems with box lengths ranging from 115 to 135 nm; specifically, 115 nm (60 mg/mL), 120 nm (52.8 mg/mL), 125 nm (46.7 mg/mL), 130 nm (41.5 mg/mL), 132 nm (39.6 mg/mL), and 135 nm (37.1 mg/mL), and performed long-timescale simulations for each. Across these systems, vesicle formation is observed under nearly all conditions, demonstrating the robustness of this self-assembled architecture (Fig. S3). The rate of vesicle formation shows a clear dependence on density. Higher-density systems consistently exhibit faster coalescence events, rapidly bringing RNA and protein chains together into a single large condensate that subsequently reorganizes into a vesicular morphology. In contrast, lower-density systems exhibit sequential assembly states, suggesting a more gradual, hierarchical pathway toward vesicle formation.

To quantify density-dependent effects, we analyze the temporal evolution of the number of clusters, the largest cluster size, and the radius of gyration of the largest cluster across all systems (Fig. 8B and Fig S4A). At higher densities, these metrics exhibit rapid convergence: the number of clusters drops sharply as chains quickly merge into a dominant aggregate, whose radius of gyration stabilizes before reorganizing into a hollow vesicle-like structure.

**Figure 8:**
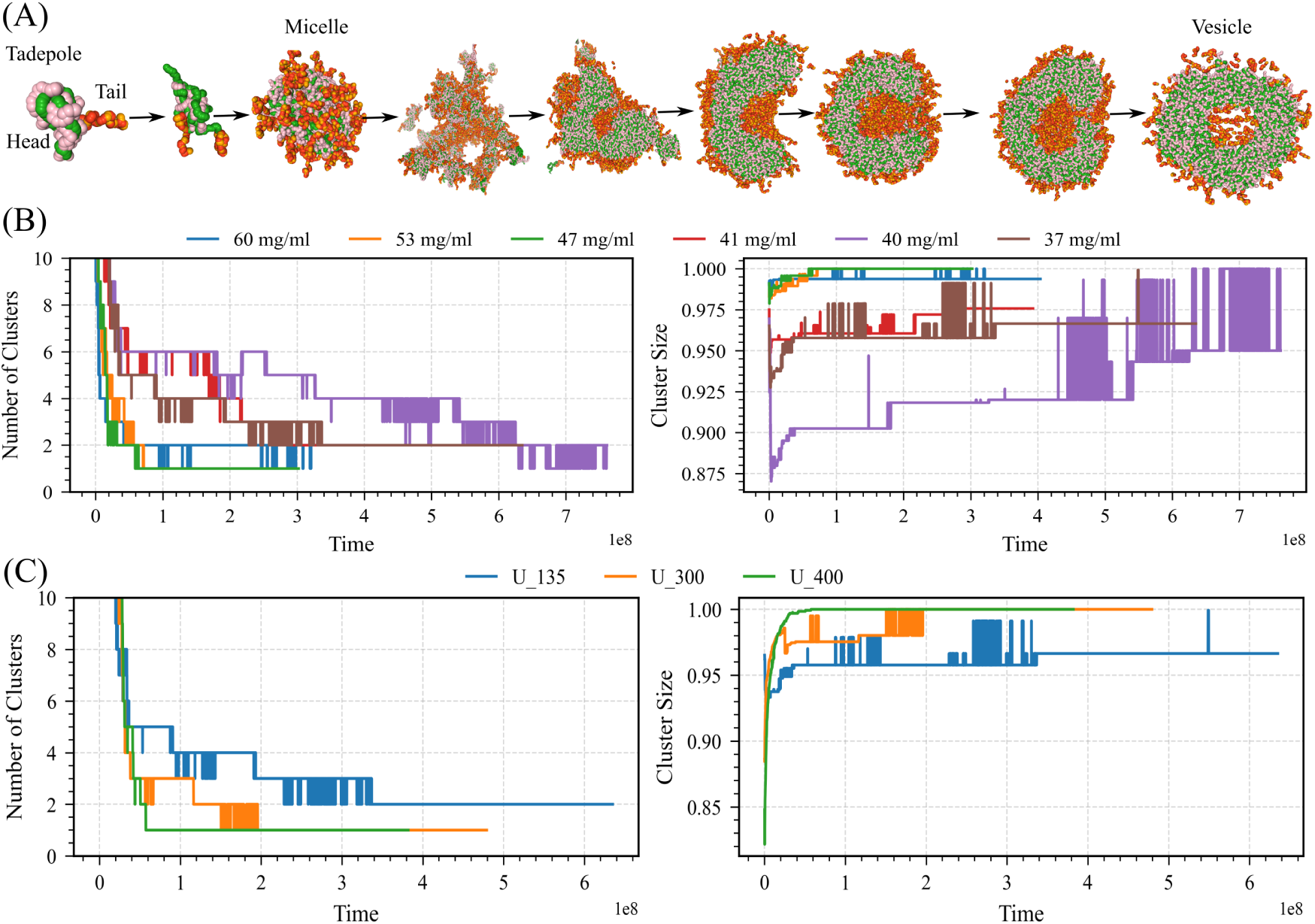
Temporal evolution of vesicle formation in the R_135_ + (N_3_G_2_)_13_ system with U_135_. (A) Representative simulation snapshots depicting the sequential structural transitions observed over time (time unit = 10 fs). The system initially forms tadpole-like morphologies, followed by tadpoles with double tails, micelles, and percolated network-like structures. These intermediates subsequently reorganize into partially opened vesicle-like assemblies and eventually mature into fully developed vesicles. (B) Quantification of density-dependent effects through the temporal evolution of key structural metrics, including the number of clusters and the cluster size distribution of the largest cluster across all systems. (C) Effect of chain length on biomolecular condensate formation: quantification of length-dependent effects based on the temporal evolution of key structural metrics, including the number of clusters and the cluster size distribution of the largest cluster across all systems.

At lower densities, RNA molecules preferentially associate with the arginine-rich peptides (Domain A), generating tadpole-like complexes rather than immediately forming a cohesive droplet. These elongated tadpole complexes subsequently coalesce into micelle-like assemblies, which later merge into a percolated network. Over time, these networks evolve into condensates containing highly open, cavity-like regions, which progressively reorganize and close to yield a fully formed multiphase vesicle (Fig. 8A and Fig. S3).

Additionally, to further investigate the role of chain length in LLPS and vesicle formation, we examine the same model system of R_135_(N_3_G_2_)_13_, each simulated with three different effective RNA chain lengths: U_135_, U_300_, and U_400_, while keeping the total number of RNA residues constant in a 135 nm box (37.1 mg/mL).

Our results show that increasing RNA chain length strongly promotes condensation. Longer chains reduce the conformational entropy penalty associated with binding and enhance multivalent interactions, thereby driving faster nucleation and accelerating the progression toward vesicular structures (Fig. 8C and Figure S4B). To quantify this effect, we analyze the temporal evolution of the number of clusters, the cluster size, and the radius of gyration of the largest cluster across all chain-length conditions. As chain length increases, the number of clusters decreases more rapidly, indicating earlier coalescence into a single dominant condensate (Fig. 8C and Fig. S4B). Correspondingly, the size of the largest cluster grows more quickly, reflecting faster assembly and structural maturation. Overall, our findings demonstrate that both higher density and longer chain length significantly accelerate condensate formation, ultimately promoting more rapid and robust vesicle formation.

## Discussions

Multiphasic condensates arise when distinct classes of intermolecular interactions drive components to partially demix rather than form compositionally uniform droplets. Such compositional structuring, including biphasic, multilayered, or vesicle-like organization, is increasingly recognized as a key determinant of functional specialization in membraneless organelles.^48–50^ Protein–RNA mixtures are particularly rich examples, yet the physical rules that map sequence-encoded interaction patterns and stoichiometry onto specific internal architectures remain incompletely understood. Using a minimal two-domain protein model and coarse-grained simulations, we identify simple principles governing the formation and stability of multiphase condensates across a broad range of interaction strengths and compositions.

Our simulations examine three interaction regimes that differ in the relative strengths of protein–RNA and protein–protein interactions. In System A, both Domain L–L and Domain L–RNA interactions are weak or repulsive, while Domain H–RNA affinity remains strong. This regime yields robustly multiphase vesicles composed of an empty core, a Domain L-rich inner layer, a Domain H–RNA intermediate layer, and a Domain L-rich outer shell. These vesicles form spontaneously from well-mixed initial conditions and do not require strong charge imbalance or influx-driven oversaturation, distinguishing them from previously studied hollow condensates. Their stability arises from a balance between strong Domain H– RNA binding and weak cohesion among domain L residues, and they appear only within a specific stoichiometric window in which this balance is maintained.

In System B, moderate Domain L–L cohesion is introduced while Domain L–RNA interactions remain weak. This change shifts the dominant architecture toward multilayered droplets with a Domain L-enriched core, an intermediate Domain H–RNA layer, and a secondary Domain L-rich outer shell. Increasing the fraction of Domain H leads to systematic structural transitions: partially covered multilayers become fully developed multilayers, which give way to dispersed Domain L-rich microdomains within an Domain H–RNA network, and eventually nearly homogeneous Domain L–RNA–dominated droplets. These transitions reflect a balance between Domain L–L attraction, which favors a cohesive core, and Domain H–RNA interactions, which drive compositional rearrangement around that core.

In System C, strong Domain L–L interactions dominate over moderate Domain L–RNA affinity, while Domain H–RNA binding remains strongest. Under these conditions, the condensates adopt biphasic architectures in which a dense Domain L-rich core is surrounded by an Domain H–RNA shell. Increasing the abundance of Domain H gradually disrupts the compact core, first allowing partial mixing and then yielding scattered Domain L-rich inclusions embedded within a continuous Domain H–RNA phase. System C thus mirrors the progression observed in System B, but the layer ordering is reversed due to the much stronger cohesion among Domain L residues.

Across Systems A–C, varying the stoichiometric ratio of Domains H and L modulates the internal organization in predictable ways. For each system, the stoichiometries that yield vesicles or well-defined multilayers coincide with minima in the PMF-weighted interaction energy, indicating that these morphologies are favored by the overall balance of effective pairwise interactions rather than by charge alone. The observed transitions, from micelles to incomplete vesicles to closed vesicles in System A, or from layered structures to fragmented microdomains in Systems B and C, are consistent with classical behavior in multicomponent polymer–amphiphilic systems but arise here solely from competing protein–RNA interactions.

We also investigated the influence of protein–RNA density and RNA chain length. Increasing density accelerates chain coalescence and leads to faster vesicle closure, whereas lower densities reveal sequential intermediates, such as tadpoles, micelles, and network-like aggregates. Similarly, increasing RNA chain length enhances multivalent binding and speeds the emergence of the final condensate architecture. Importantly, these perturbations influence kinetics but not the ultimate structure: the vesicle, multilayered, or biphasic architectures are robust to changes in density and chain length, further highlighting that morphology is governed primarily by interaction grammar and stoichiometry.

Together, these findings suggest three general design principles. First, when heterotypic Domain H–RNA binding strongly dominates and auxiliary interactions among Domain L residues remain weak, vesicle-like multiphasic condensates emerge naturally without requiring charge asymmetry. Second, moderate Domain L–L cohesion favors multilayered droplets whose organization depends sensitively on the Domain H:L stoichiometric ratio. Third, strong Domain L–L cohesion promotes biphasic structures with dense cores that are progressively destabilized as Domain H becomes more abundant. These principles unify observations across all three systems and provide a framework for predicting how specific sequence patterns and compositions shape the architecture of condensates.

The structural motifs identified here connect naturally to experimental observations. Reconstituted protein–RNA systems frequently display competitive binding, demixing, and layered organization, and our results suggest how specific sequence alterations, such as increasing aromatic content in Domain L or modifying charge patterning, could shift condensates between vesicular, multilayered, or biphasic states. The prediction that vesicles can form spontaneously under charge-balanced conditions is particularly notable, as most reported hollow condensates arise from influx-driven or oversaturated states.^21^ Matching the predicted stoichiometric windows or manipulating RNA chain length offers straightforward routes for experimental validation.

Beyond biological relevance, the vesicle-like structures observed here resemble amphiphilic assemblies despite being composed entirely of intrinsically disordered protein segments and RNA.^51,52^ This points to opportunities for engineering membrane-free vesicles with programmable permeability, encapsulation, and responsiveness, with potential applications in synthetic organelles, microreactors, and stimulus-controlled delivery systems.^53,54,54–57^

Overall, our study provides a mechanistic map linking sequence grammar, stoichiometry, and intermolecular competition to the emergence of multiphasic protein–RNA condensates. The simplicity and generality of these principles make them well-suited for guiding future *in vitro* reconstitution, for interpreting the internal organization of cellular condensates, and for designing programmable biomolecular materials.

## Acknowledgments

This work was supported by funds from the National Institute of General Medical Sciences, with grant no. R35 GM138243 awarded to DAP.

## TOC Graphic

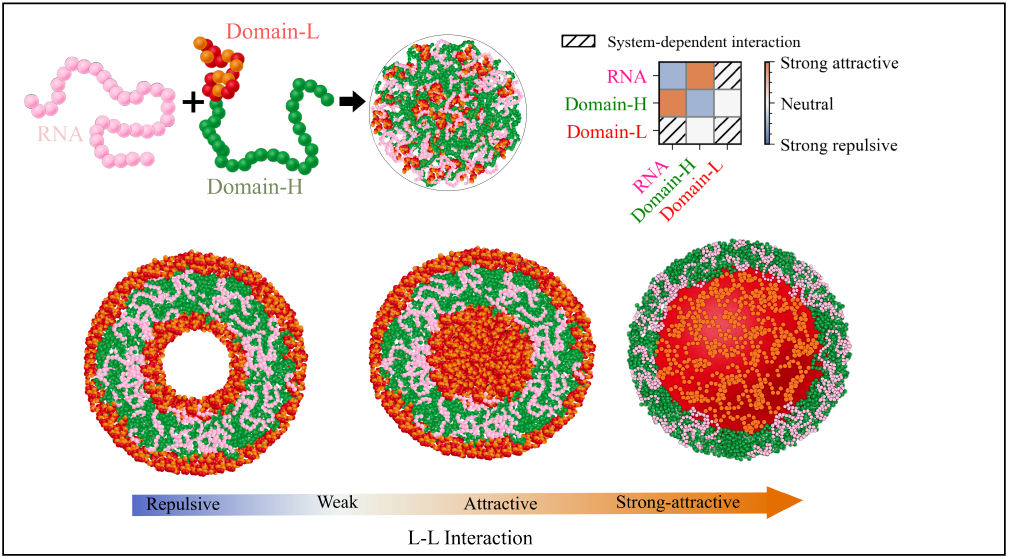

